# Why are telomeres the length that they are? Insight from a phylogenetic comparative analysis

**DOI:** 10.1101/2024.09.25.613781

**Authors:** Derek M. Benson, Dylan J. Padilla Perez, Dale F. DeNardo

## Abstract

Telomeres are short repeating nucleotide sequences at the ends of chromosomes that shorten with every cellular replication. Despite the importance of keeping telomere length within a critical homeostatic range, adult telomere length can differ by two orders of magnitude across vertebrate species. Why telomere length varies so widely remains unknown, though popular hypotheses suggest that body size, lifespan, and endothermy are key variables that have coevolved with telomere length. To test the relationship among telomere length, telomerase activity (which extends telomeres), and these variables, we modeled the evolution of telomere length across 122 vertebrate species. We failed to find an influence of body mass, lifespan, or baseline metabolism on telomere length. However, we found a significant interactive effect between baseline metabolism and body mass. The presence of telomerase activity was positively correlated with telomere length across the 58 species where data for both existed. Taken together, our findings suggest that body mass may have differentially influenced the evolution of telomere length in endotherms and ectotherms and indicate that telomerase activity and telomere length may have coevolved.

## Introduction

Telomeres are DNA sequences that are critical for protecting the ends of chromosomes. Accordingly, all vertebrate chromosomes end in telomeres. Yet, despite the crucial role of telomeres and their ubiquity across vertebrates, the existing literature on absolute telomere length has not previously been comparatively analyzed to better understand the variation in telomere biology across major vertebrate clades.

Structurally, vertebrate telomeres are comprised of repeating TTAGGG nucleotide sequences and a protein complex known as shelterin (Rhodes and Giraldo, 1995; De Lange, 2005). The 3’ end of telomeres ends in a G-strand overhand, which often forms a T-loop structure by invading and binding to double-stranded telomeric DNA (Cong et al., 2002; Haussmann and Marchetto, 2010). The functional limitations of DNA polymerase and the creation of Okazaki fragments lead to the progressive loss of telomeric DNA with each cellular replication (Gilson and Géli, 2007). Many species, including humans, experience replicative aging where cells enter senescence and stop replicating when telomere length hits a critical minimum (Saretzki and Von Zglinicki, 2002; Shay and Wright, 2005). In addition to the impacts of cellular replication, oxidative stress can also cause significant telomere shortening (Von Zglinicki, 2002). The telomeric 5’-GGG-3’ sequence is particularly susceptible to damage by reactive oxygen species and conversion of one or more guanines into 8-oxoG (Kawanishi and Oikawa, 2004). Furthermore, the DNA damage response is repressed at telomeric DNA, and telomeric DNA within the single-stranded T-loop is not protected by hydrogen bonding and, due to the absence of a template strand, not able to be repaired by base excision repair (Ahmed and Lingner, 2018). In contrast to the cellular senescence associated with short telomeres, long telomeres are associated with increased cancer risk in some species, so it is crucial for an organism to maintain its telomere lengths within a critical range (Rode et al., 2016; Aviv et al., 2017; McNally et al., 2019).

To counter the progressive shortening of telomeres resulting from cellular replication, organisms can prevent telomere shortening through antioxidant production and, in some cases, elongate telomeres via telomerase activity and alternative telomere lengthening (Neumann et al. 2013; Armstrong and Boonekamp 2023). Telomerase is a ribonucleoprotein reverse transcriptase comprised of catalytic (TERT) and RNA (TERC) components (Cong et al., 2002). Telomerase extends the G strand of telomeric DNA, after which the C strand is filled in by DNA polymerase α *− primase* (Bonetti et al., 2014). Despite its obvious benefits, in humans and many endotherms telomerase activity is suppressed in most somatic tissues outside of embryonic stages (Wright et al., 1996). It has been hypothesized that this inactivity of telomerase is an anti-cancer protection mechanism, as telomerase activity enables continuous cellular replication and is almost always necessary for cellular immortalization (Shay and Wright, 2011). In fact, telomerase activity is present in 85-90% of all human tumors (Shay and Bacchetti, 1997).

Since their discovery, telomeres, telomere length, and telomerase activity, collectively referred to as “telomere dynamics,” have been studied extensively for their potential involvement in both aging and cancer (Aubert and Lansdorp, 2008; Xu et al., 2013). Additionally, many studies have investigated telomere dynamics in non-human vertebrates, revealing that telomere length, replicative senescence, and repressed telomerase activity are not conserved traits but rather vary among species (Gomes et al., 2010). In fact, telomere length ranges across species from one to hundreds of kb (Rocco et al., 2001; Zhdanova et al., 2005). Why telomere length varies so widely across vertebrates remains to be understood, though popular hypotheses suggest that body size, lifespan, and endothermy are key variables that have coevolved with telomere length (Heidinger et al., 2012; Olsson et al., 2018; Pepke and Eisenberg, 2022). Larger body size increases the number of cells an organism has which increases the number of potentially cancerous cells, increased lifespan enables more mutations to accumulate over time, and endothermy increases the rate of metabolism and cellular turnover, thus increasing the mutation rate of cells (Peto et al., 1975; Seluanov et al., 2008; Hayes, 2010).

While the importance of telomere dynamics has resulted in a considerable number of publications regarding the subject across numerous species, no literature has taken a comprehensive, wide-scale comparative approach to review and analyze the relationships among telomere dynamics, body size, lifespan, and endothermy across major vertebrate clades. The goals of this review are to 1. Determine whether lifespan, body mass, and endothermy have affected the evolution of telomere length; 2. Elucidate whether telomere length is differentially affected by these variables among major vertebrate clades; and 3. Investigate whether telomere length and telomerase expression have coevolved.

## Materials and Methods

### Literature search and data collection

To search for existing literature on telomere length and telomerase activity, key terms including “telomere length”, “telomerase activity”, and specific species or taxonomic groups were input into both Google Scholar and Web of Science (Figure S1). Given the discrepancies in telomere length measurements across methodology, we only collected data from studies using telomere restriction fragment analysis (TRF) as it is the most widely used quantitative measurement of telomere length. As TRF can be performed with or without completely denaturing the DNA and measuring interstitial telomere sequences (ITSs), we tested for an effect of denaturing on telomere length across all the species in our analyses. As we did not detect any significant effect, we compared telomere length across the methodologies, though we cannot completely discount the fact that methodology may influence our findings. When available and for consistency, we utilized average adult telomere length as reported in previous analyses that had collected data across multiple studies (see, Gomes et al., 2011; Criscuolo et al., 2021; Pepke and Eisenberg, 2022). For species not included in these analyses, we used the average telomere length value reported in individual studies. For studies that did not report an average telomere length, we calculated average telomere length as the mean telomere length across all individuals in a study. We did not include studies or datapoints on telomere length in extremely young or old individuals (see the supporting information for more information on the collection of data). Given that telomere length is generally strongly correlated but not constant across somatic tissues within individuals (McLester-Davis et al., 2023), we tested for an effect of tissue type on telomere length. As we did not detect an effect of tissue type on telomere length, we compared telomere length across tissue types (see supporting information for more detail). Additionally, as previous studies have shown no evidence for sex-based differences in adult telomere length across vertebrates, we included data for both sexes in our analyses (Remot et al., 2020). Telomerase activity in the analyzed cells (hereafter referred to as telomerase activity) was recorded as a binary variable, being either absent or present in the somatic tissue of the study species. Telomerase activity data from gonadal tissue were not utilized as many species exhibit telomerase activity in gonadal tissue but not in other somatic tissues (Wright et al., 1996; Sun et al., 2021). For studies that reported telomere length but not telomerase activity, we recorded telomerase activity as not available (n/a) for the species. As telomere length data from Seluanov et al. (2007) was found to be highly correlated with but longer than that of overlapping species analyzed by Gomes et al. (2011), we applied the same correction factor (0.6617 x TL – 2.4454 kb) that was utilized by Pepke and Eisenberg (2022). Once telomere length and telomerase activity data were collected, we obtained species-specific maximum lifespan in years (hereafter referred to as lifespan) and adult mass in grams primarily from the AnAge database (de Magalhães et al., 2024; https://genomics.senescence.info/species/index.html). For species that did not have an entry in the AnAge database, information on adult mass and lifespan was collected from various literature sources (see supplemental table 1 for more information). If no data on lifespan or adult mass could be found, the species was not used for any analysis. Additionally, as domestication may have influenced the evolution of telomere length, animals with a long history of domestication were not included in our analyses. We utilized telomere dynamics data for 122 vertebrate species across 42 orders and 88 families.

### Vertebrate phylogeny

To build a vertebrate phylogeny for this review, we used the Build a Timetree function available at The TimeTree of Life database (https://timetree.org/), which outputs a timetree of taxa of interest extracted from the global timetree connecting species and publication-specific timetrees in the database. For species that did not have phylogenetic data readily available, we utilized closely related species or congeners when constructing the phylogeny (see the supporting information for more detail). Although the placement of taxa in our species-level tree of vertebrates was generally well supported, phylogenetic comparative methods appear to be robust to perturbations such as tree misspecification or soft polytomies (Stone, 2011).

### Statistical Analysis

To better understand the evolution of telomere length, we estimated its ancestral state across vertebrates. We used the function *fastAnc* in the “phytools” package of R v. 4.2.2 (Revell, 2022), which performs a reasonably fast estimation of the most likely ancestral states for a continuous trait by taking advantage of the fact that the state computed for the root node of the tree during Felsenstein (1985) contrasts algorithm is also the most likely estimate of the root node. Thus, the function re-roots the tree at all internal nodes and computes the contrasts state at the root each time. The function also computes variances or 95% confidence intervals on the estimates.

Subsequently, we used Phylogenetic Path Analysis (van der Bijl, 2018) to test our hypotheses on the relationships among body mass, baseline metabolism (i.e., the dichotomous distinction between ectotherms and endotherms), lifespan, and telomere length across species. Phylogenetic path analysis uses Phylogenetic Generalized Least-squares (PGLS) to perform confirmatory path analysis based on the directed separation method–“d-separation”–as described in Shipley (2013). We examined the explanatory power of 8 competing models involving different relationships among variables, which enabled us to identify the most plausible hypothesis given the available data, and thus infer the relative importance of mechanisms described by optimality models. To select the most likely path model, we relied on the *C* statistic of the model, which is a maximum likelihood estimate that follows a x^2^ distribution with degrees of freedom df = 2k. Therefore, it provides a convenient statistic for testing the goodness of fit of the whole path model (Shipley, 2013). If the path model did not provide a good fit to the data because the p-value of the *C* statistic was below the alpha value (i.e., < 0.05), we proceed to examine the conditional independencies tested in the path model. A conditional independency specifies the list of pairs of variables that are statistically independent conditioning on a set of other variables in the causal path model. A conditional independency is only supported by the data if its p-value is greater than the alpha value (i.e., p-value > 0.05). The most likely conditional independency suggested by path analysis was then refitted assuming that the species trait values evolved via a Brownian motion model, an Ornstein–Uhlenbeck motion model, and a Pagel’s lambda model using the R package ‘ape’ v. 5.6.2. To evaluate the models’ goodness of fit, we used information-theoretic approach such as AICc. We ranked the candidate models accordingly and selected the most likely one for inferences (lowest value of AICc). To investigate the effects of sample size on the exploratory power of our models, we conducted a sensitivity analysis across the entire dataset manually utilizing the previously determined most likely model. In brief, we ran 1000 simulations for each of five bins randomly removing 10, 20, 30, 40, and 50% of species, for a total of 5000 simulations. For each bin, we calculated the mean parameter estimate for the model intercept, body mass, lifespan, baseline metabolism, and the baseline metabolism x body mass interaction. Additionally, we calculated the standard deviation for each parameter, and the 95% CI for each parameter. Significant parameter estimates were those where the 95% CI did not include 0.

We also estimated the degree of phylogenetic signal in our data based on the most likely model. The “ape” package of R enabled us to modify the correlation structure of the residuals in the function gls. Thus, we assigned different fixed values to the parameter λ, ranging from zero (no phylogenetic signal, equivalent to a “star” phylogeny) to one (consistent with Brownian motion). Intermediate values of λ imply that the data support a model that is somewhere between a “star” phylogeny and Brownian motion. λ represents the amount of phylogenetic “signal” in the data, conditional on the tree. In a regression context, we used maximum likelihood to simultaneously fit the regression model and estimate λ. Because confidence intervals for λ were provided by the models, we tested the hypotheses that λ = 0 (i.e., independence) and λ = 1 (Brownian Motion).

Lastly, we investigated whether telomerase activity changes as a function of telomere length. To do that, we modeled the correlated evolution of telomere length and telomerase activity under the threshold model (Felsenstein, 2012). This model creates a natural framework by which one can evaluate the evolutionary correlation between a continuous variable (e.g., telomere length) and a discrete variable (e.g., telomerase activity). In R, the correlational threshold model can be performed through the function threshBayes available in the package “phytools” (Revell, 2022). It also enabled us to generate a three-panel plot showing the likelihood profile, the mean acceptance rates (using a default sliding window), and a profile plot for the correlation coefficient (Figure S2). In addition, we used the density function to later plot a posterior density of the correlation coefficient and added a 95% high probability density interval (HPD) around the value of *r*. We did this by extracting a post-burn-in sample of values of *r* (assuming a 20% burn-in) and using the function HPD interval from the package “coda”. To produce good visualization of our results and ensure that they are reproducible, we carried out all the analyses in the free software for statistical computing R v.4.2.2 (R Core Team, 2022).

## Results

Our results indicated no effect of lifespan on telomere length. However, the evolution of telomere length across vertebrates appeared to be influenced by the interaction between baseline metabolism and body mass of species (Figure S3-4). The information-theoretic approach revealed that a model including the main effects of these variables was strongly supported (Table 1). Generally, telomere length increased with body mass in ectotherms and decreased with body mass in endotherms (Figure 2). Finally, our analyses indicate that there may have been many transitions in telomere length across vertebrates (Figure 1).

**Table 1:** Contrast of parameters estimated by the most likely model according to the information-theoretic criteria of selection.

|  | t-value | p-value |
| --- | --- | --- |
| (Intercept) | 6.84 | <0.001 |
| Baseline metabolism | 0.81 | 0.42 |
| Log mass | 1.11 | 0.27 |
| Log lifespan | 1.37 | 0.18 |
| Log mass x Baseline metabolism | 2.41 | 0.018 |

**Figure 1:**
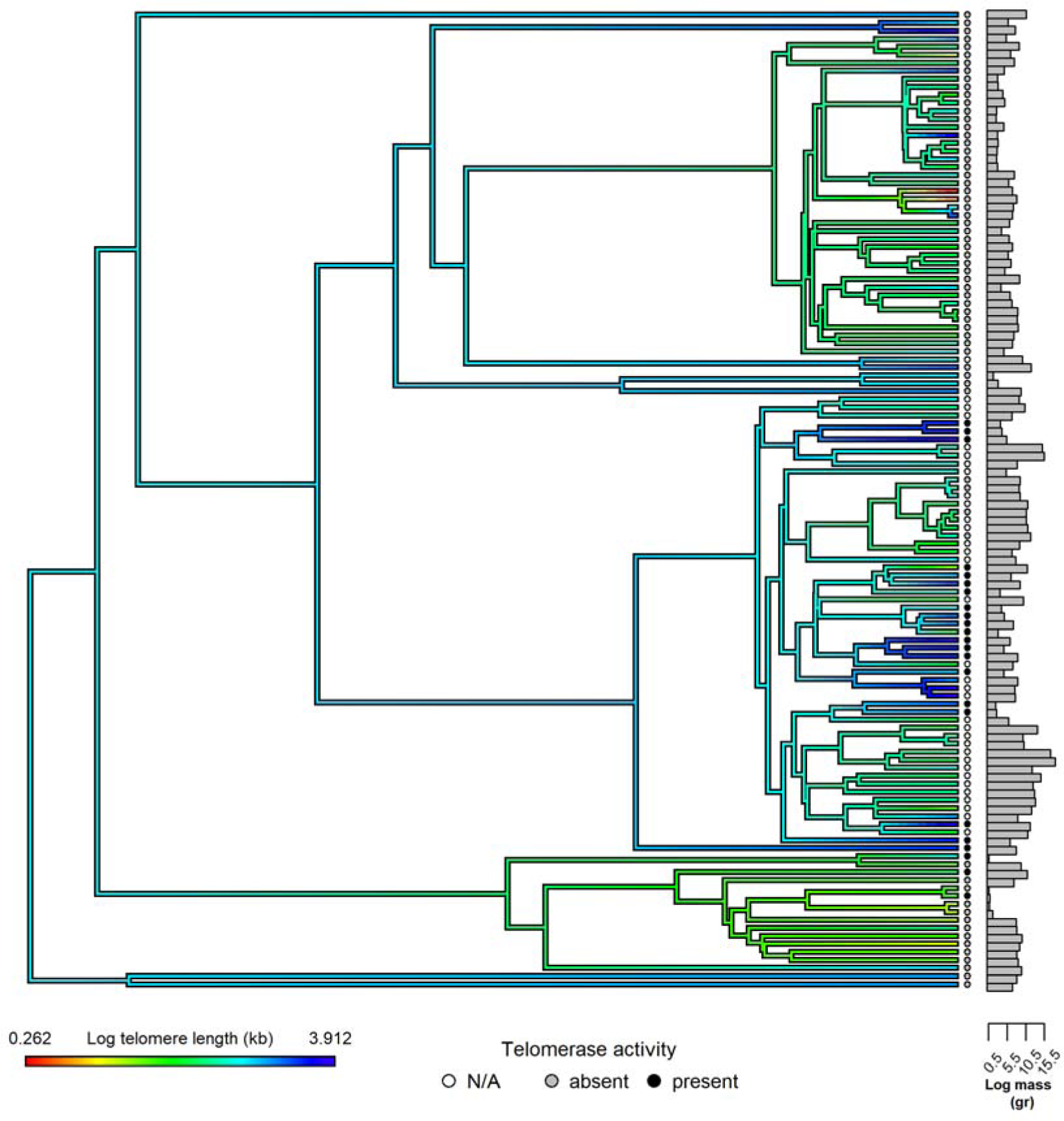
Telomere length, body mass, baseline metabolism, and telomerase activity across 122 vertebrate species. Warmer colors represent shorter telomeres.

**Figure 2:**
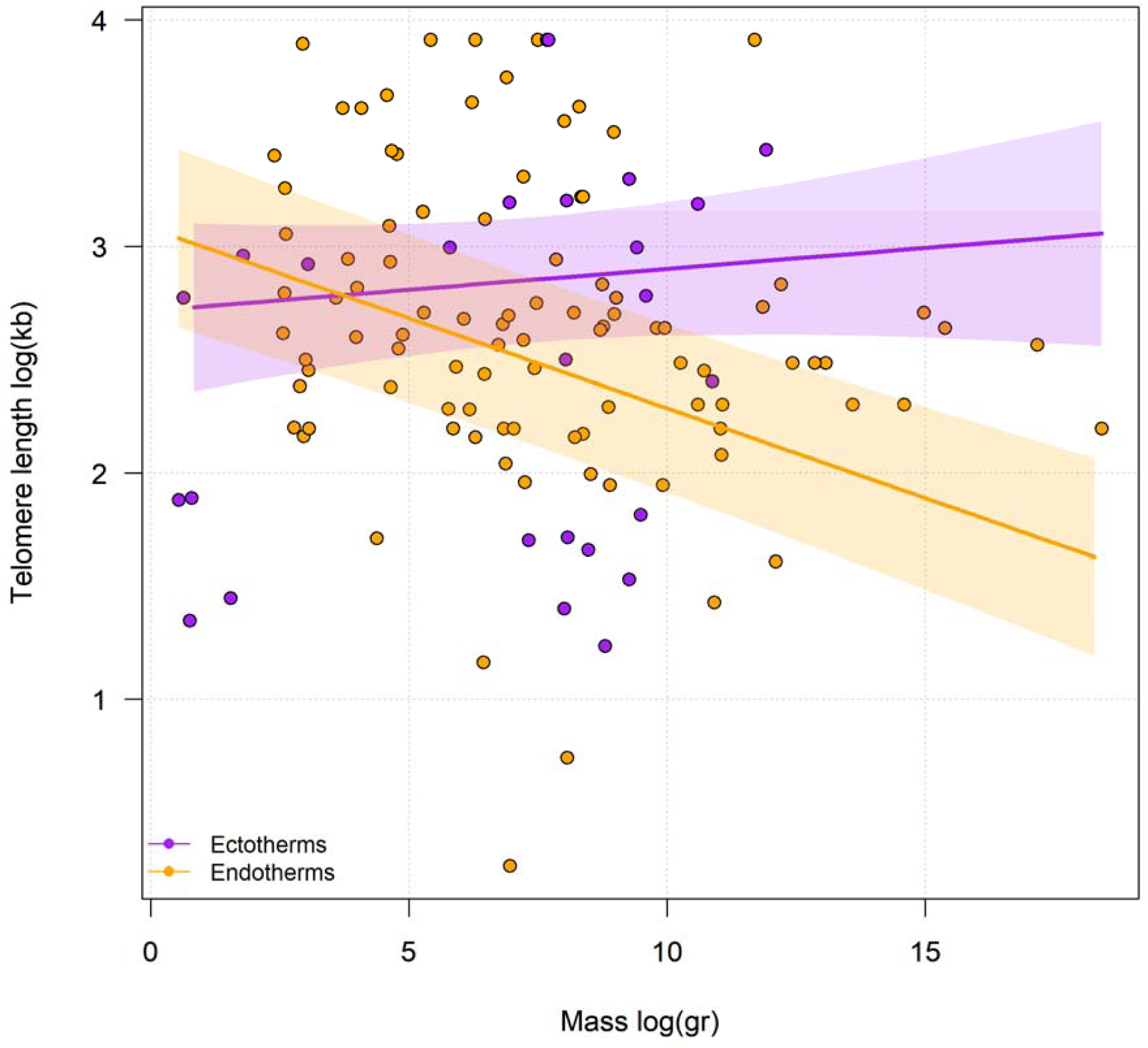
Evolution of telomere length as a function of the interaction between body mase and baseline metabolism across vertebrates. Shading around the regression lines represents the standard error.

As the scatter in the data indicates substantial noise, we examined the impact that different sample sizes could have on our results. Randomly removing 10, 20, 30, 40, and 50% of species in the analyses did not affect the likelihood of finding a significant effect of the interaction between baseline metabolism and body mass on telomere length, though the independent effects of lifespan, baseline metabolism, and body mass were significantly affected by sample size (Figure S5).

Based on a likelihood ratio test (X2), models with λ fixed at 0 and 1 were significantly different from one that had λ fixed at 0.5. Thus, we rejected the hypotheses of independent residuals and Brownian motion. The most likely model suggested an intermediate degree of phylogenetic signal in the data.

Finally, the threshold model of correlated evolution indicated that the evolution of telomere length is correlated with the presence of telomerase activity. According to the posterior probability distribution of the correlation coefficient (r = 0.51), the presence of telomerase activity significantly increased with telomere length. This conclusion was based on the 95% high probability density (HPD = 0.18 − 0.79) interval around *r*, which did not include 0 (Figure 3 and S2).

**Figure 3:**
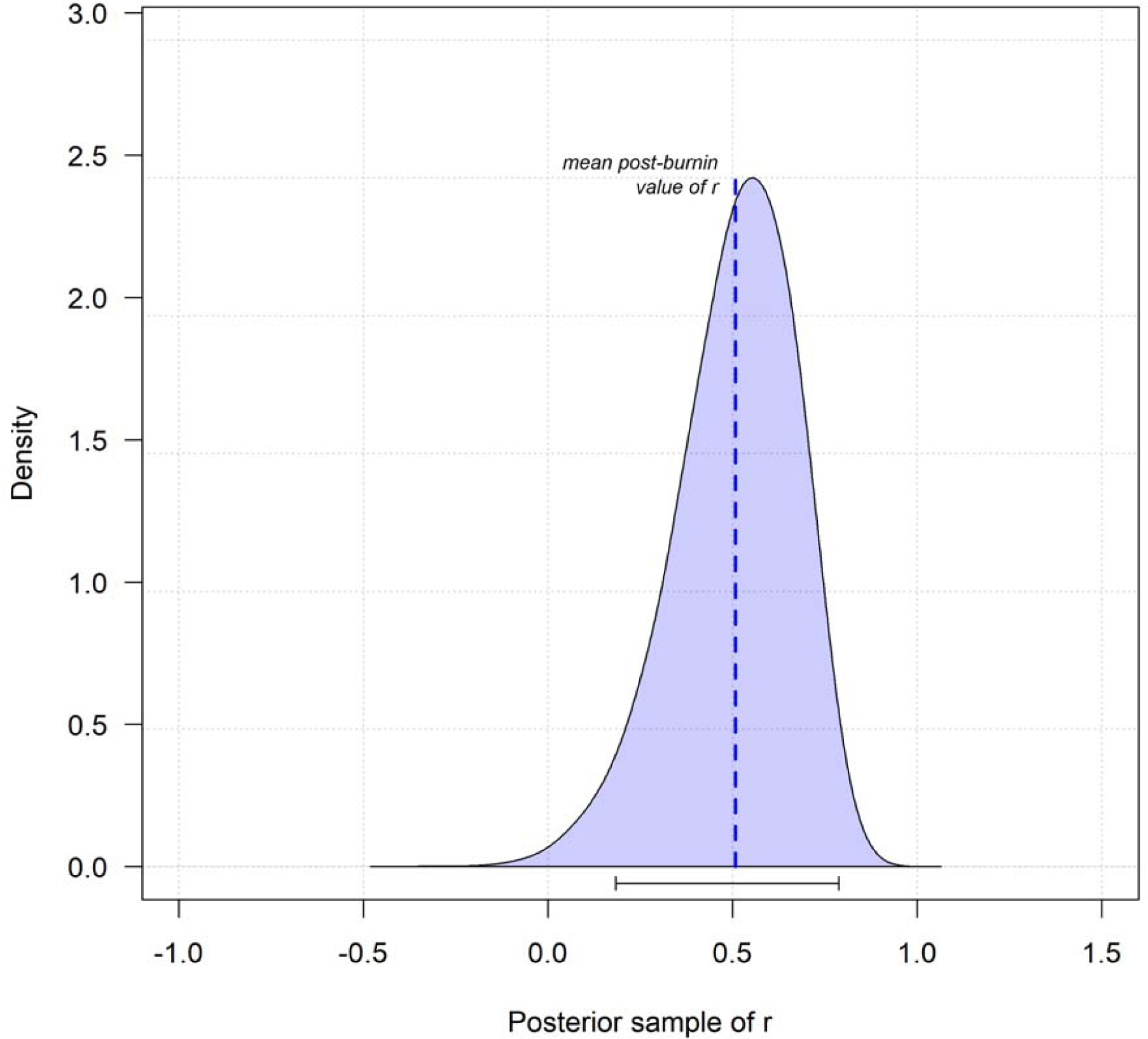
Posterior density of the correlation between telomere length and telomerase activity in vertebrates. The dashed line represents the estimated correlation coefficient (r = 0.51), representing the correlation between telomere length and the presence or absence of telomerase activity. Additionally, the 95% high probability density (HPD = 0.18 − 0.79) is represented by the solid black bar below the distribution.

## Discussion

To our knowledge, ours is the first study to test for correlates between telomere length and various variables across vertebrate clades, though previous studies have investigated within clades (Gomes et al., 2011; Criscuolo et al., 2021; Pepke and Eisenberg, 2022). Across vertebrates, we found evidence that the interaction between baseline metabolism and body mass is correlated with telomere length. However, we failed to find evidence that body mass, lifespan, baseline metabolism, or any other interactions are correlated with telomere length across major vertebrate clades. It should be noted that while we did not detect an effect of tissue type or methodology for our analyses, we cannot discount the possibility that the heterogenous nature of the data analyzed may influence our results and represent a major limitation of our study. Still, our study lends valuable insight into the evolution of telomere length and suggests that the evolution of telomere length has been influenced by baseline metabolism and that the effect of body mass on telomere length may differ between endotherms and ectotherms.

The relationship between body mass and telomere length remains unclear, as previous research on birds and mammals has yielded contrasting results (Gomes et al. 2011; Criscuolo et al. 2021; Pepke and Eisenberg 2022). As we included fishes and ectothermic diapsids, our results are not directly comparable, though they suggest that the correlation between body mass and telomere length may be dependent on baseline metabolism. One possible explanation for the differential effect of body mass based on baseline metabolism is that high stable body temperatures may be associated with an increased risk of malignant tumors (Olsson et al. 2018). While plausible, many ectotherms prefer and maintain body temperatures above that of endotherms, and previous research has failed to find a relationship between high stable body temperature and an increase in malignant tumors (Stevenson 1985; Olsson et al. 2018). Alternatively, it may be that the difference in metabolic rate between endotherms and ectotherms has significantly affected the evolution of telomere length, as metabolic rate has been shown to be positively correlated with DNA damage (Moretton and Loizou 2020). Thus, the selective pressure that body mass imposes on telomere length may differ between endotherms and ectotherms, as is evident by the difference in slopes relating telomere length to body mass. Regardless of the reason behind the relationship between baseline metabolism and telomere length, it is important to consider the disparity in sample size between endotherms and ectotherms in our analyses. Vertebrate ectotherms are understudied in terms of telomere biology when compared to endotherms, and information on major clades including Anurans and Agnatha is nearly or completely absent. Given the importance of these groups for understanding vertebrate evolution, it is likely that quantifying telomere length in species from these clades will further elucidate how baseline metabolism is related to telomere length. Nevertheless, our results indicate that baseline metabolism and body mass may have played a significant role in the evolution of telomere length.

In contrast to previous research in mammals (Pepke and Eisenberg 2022), we failed to find evidence that telomere length is correlated with lifespan. Conclusions on this relationship remain divided, with studies both supporting and refuting the idea that a correlation exists (Gomes et al. 2011; Tricola et al., 2018; Criscuolo et al., 2021; Pepke et al., 2023b). This discrepancy may in part be explained by the use of either early-life (within the first few weeks of life) or adult telomere length. We used adult telomere length for several reasons: Firstly, early-life telomere length changes rapidly for many though not all species (Watson et al., 2015; Pepke et al., 2023a). Thus, early-life telomere length may not matter as much as preserved telomere length. Secondly, early-life telomere length fails to account for telomere length modulation later in life. Several long-lived species have been shown to increase telomere length in adulthood (Haussmann et al., 2003). Furthermore, telomere rate of change has been repeatedly shown to be correlated with lifespan (Tricola et al., 2018; Criscuolo et al., 2021; Whittemore et al., 2019). While adult telomere length does not measure this, adult telomere length is affected by shortening and lengthening and may serve as a more effective measurement of telomere length for a species. However, among adults, age can vary considerably in longer lived species and such variation in age among adults could confound results.

Despite variation in telomere length within and among clades, our analyses indicate that long telomeres are correlated with the presence of adult telomerase activity. Long telomere length and the presence of telomerase activity might be thought to increase cancer risk as previous literature suggests that selection favors short telomeres and the repression of telomerase as a mechanism to prevent cancer (Risques and Promislow, 2018). However, while cancer is detrimental to organismal health, the vast majority of wild animals do not live to the age where cancer becomes a significant risk factor (Rozhok and DeGregori, 2016). Thus, other variables such as pace of life and reproductive investment, rather than cancer risk, may have exerted stronger selective pressures on the evolution of telomere length and telomerase activity. Additionally, given that our and previous analyses are strongly biased towards mammal and bird, it is possible that the correlations between telomerase activity, long telomeres, and cancer risk are not reflective of vertebrates in general (Gomes et al., 2011; Pepke and Eisenberg, 2022). For example, the limited data in fishes demonstrate that most species in our analysis display telomerase activity with telomeres <15 kb in length, a phenomenon rarely seen among mammals (Zhdanova et al., 2005; Elmore et al., 2008; Gomes et al., 2011).

Making comparisons across and among subclades of a larger tree can prove valuable, particularly where large timescales are involved and thus where evolutionary patterns and processes are unlikely to be constant. However, there are important considerations when choosing which clades to study, including sample size and whether there is sufficient within-clade diversity for statistical analysis (Baker et al., 2021). Given our findings and the indications that baseline metabolism may play a role in shaping the evolution of telomere length as well as the current literature’s bias towards mammals and birds, future studies need to include a wider range of vertebrates, preferably with a similar degree of representation across the major taxa, to gain a more thorough understanding of (1) how telomere length varies within and among major clades, and (2) how the effects of variables such as lifespan and body mass on telomere length may differ among clades.

Our data may suffer from both an extremely variable number of species per clade and a large variation in telomere size among species, preventing us from making a general conclusion about the evolution of telomere size across vertebrates. While sample size may have influenced our results, our findings appear to be fairly robust given that the random removal of up to 50% of the species in our study did not affect the chance of finding a significant correlation between telomere length and the interaction between body mass and baseline metabolism. In support of the idea that sample size may be impacting our findings, previous studies have found differing results depending on the clade and range of taxa used when investigating the relationship between telomere length and lifespan (Seluanov et al., 2007; Gomes et al., 2011; Criscuolo et al., 2021; Pepke and Eisenberg, 2022). For instance, work in rodents found no evidence that lifespan was correlated with telomere length, while several analyses that include a wider range of mammals have concluded that telomere length is strongly correlated with lifespan (Seluanov et al., 2007; Gomes et al., 2011; Pepke and Eisenberg, 2022).

While our greater overall sample size may explain the differences between our results and those of previous studies, many variables, such as diet and habitat, may have also significantly influenced the evolution of telomere length. While diet has been shown to significantly affect telomere length in mice and humans, it is not known whether carnivory or herbivory differentially impact telomere length in mammals (Paul, 2011; Gokarn et al., 2018). Furthermore, research on a wide range of neotropical bat species found that frugivorous, omnivorous, and carnivorous species differ in oxidative stress and antioxidant capacity (Schneeberger et al., 2014). Given that telomere length is associated with oxidative stress, it may be that diet has had a significant impact on the evolution of telomere length (Von Zglinicki, 2002). In addition, habitat and environment have been shown to impact telomere length. For example, moose (*Alces alces*) populations in boreal regions have significantly longer telomeres than do those in sarmatic mixed forest and montane regions (Fohringer et al., 2022). Furthermore, tropical and temperate birds have differentially evolved responses to oxidative damage, and telomere length in young birds of the genus *Saxicola* differs between species in tropical and temperate regions (Jimenez et al., 2013; Apfelbeck et al., 2019). In addition to needing greater and more balanced phylogenic coverage, future studies investigating the evolution of telomere length should also investigate a wider range of life history and ecological traits.

In conclusion, despite the possible complications including a mammal-biased sample size, our study provides evidence that telomerase activity and telomere length in adult life have coevolved, with longer telomeres being associated with the presence of telomerase activity. Additionally, we show that the interaction between body mass and baseline metabolism is related to telomere length. Finally, we failed to find any relationship between lifespan and telomere length. Future studies should incorporate a wider range of non-mammalian species in their analyses to further our understanding of the variables affecting telomere length while also attempting to account for other potentially confounding variables such as unbalanced sample sizes, chromosome size, diet, habitat, pace of life, and additional life history traits.

## Supporting information

Supplementary table 1 references

## Acknowledgements

We thank the anonymous reviewers for feedback on earlier drafts.

## Data Accessibility Statement

A fully reproducible workflow of the data analyses, including R scripts and additional supporting material, is available in the following repositories: Github https://dylan-padilla.github.io/telomere_evo/, a Dryad link will be available upon acceptance.

## Author Contributions

Derek M. Benson conceptualized the study, collected and curated the data, assisted with the methodology and data analysis, and wrote the manuscript. Dylan J. Padilla-Perez was responsible for the majority of the methodology and data analysis, created the figures, and assisted in writing the manuscript. Dale F. DeNardo oversaw the project and contributed to conceptualization and writing.

## Supplementary Material

**Figure S1.**
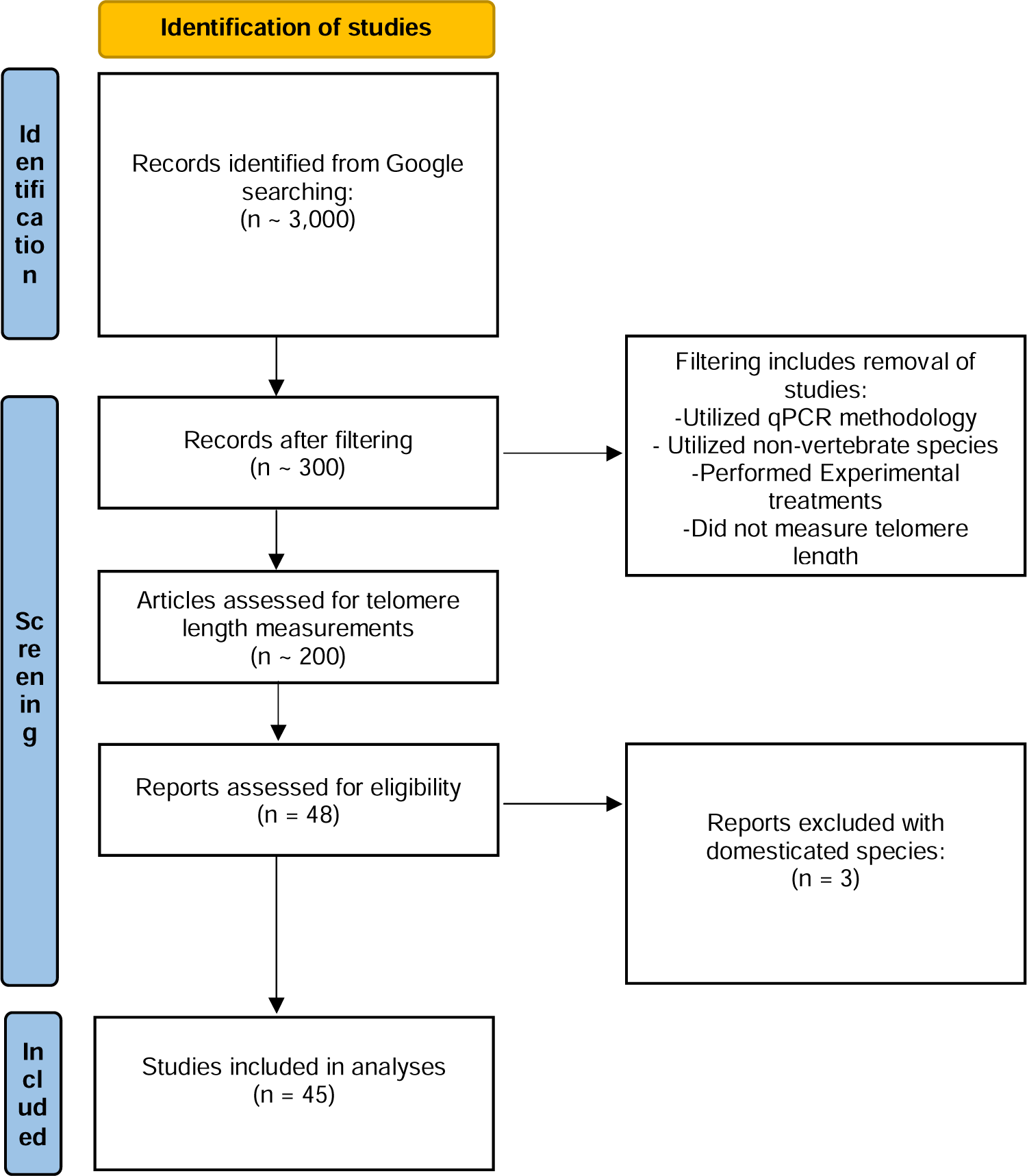
PRISMA flow diagram depicting the data collection for this study.

**Figure S2:**
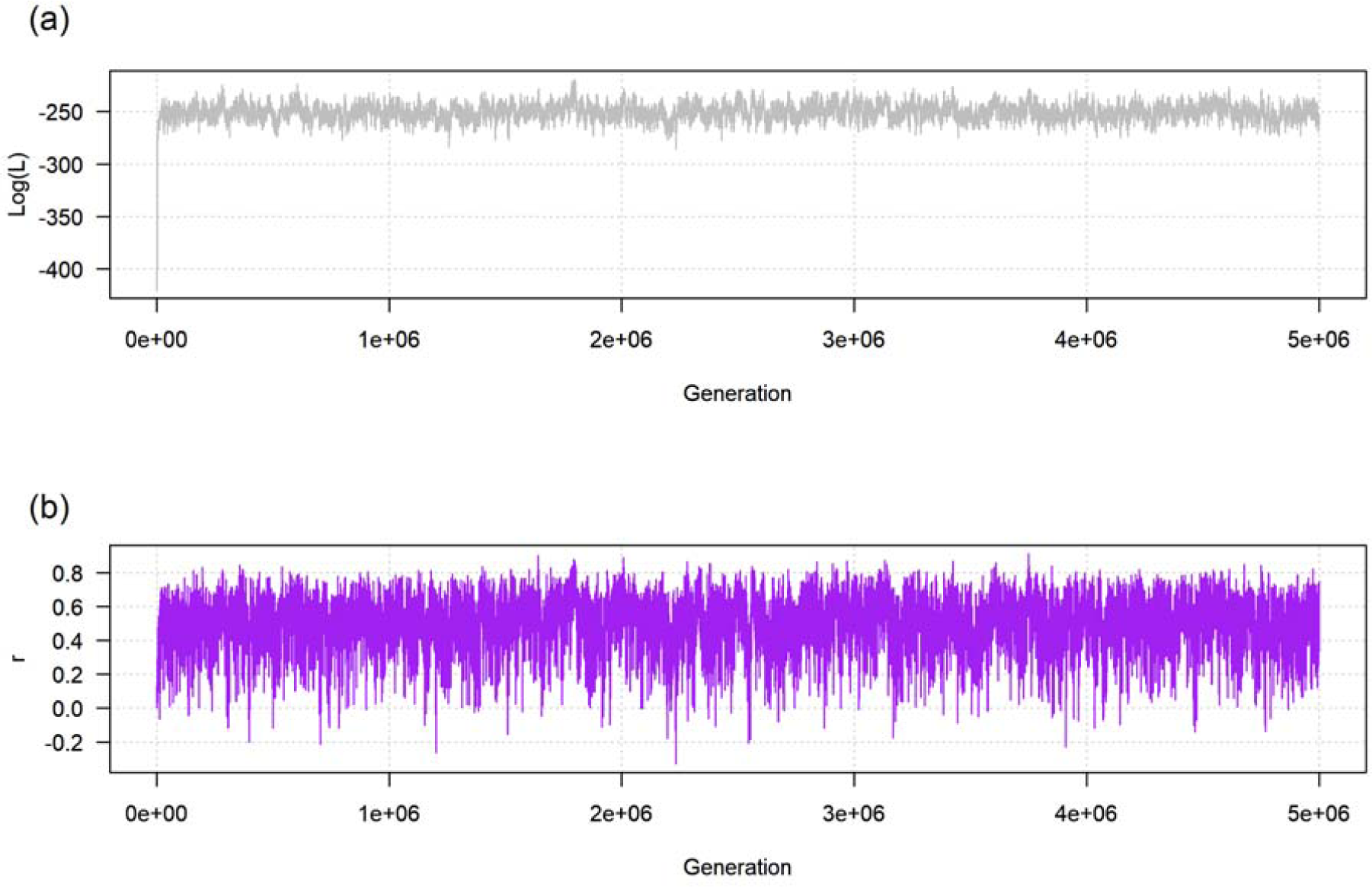
Profile plots from a Bayesian MCMC analysis of the threshold model for telomere length and telomerase activity in vertebrates. (a) The likelihood profile and (b) Profile of the correlation coefficient, r.

**Figure S3:**
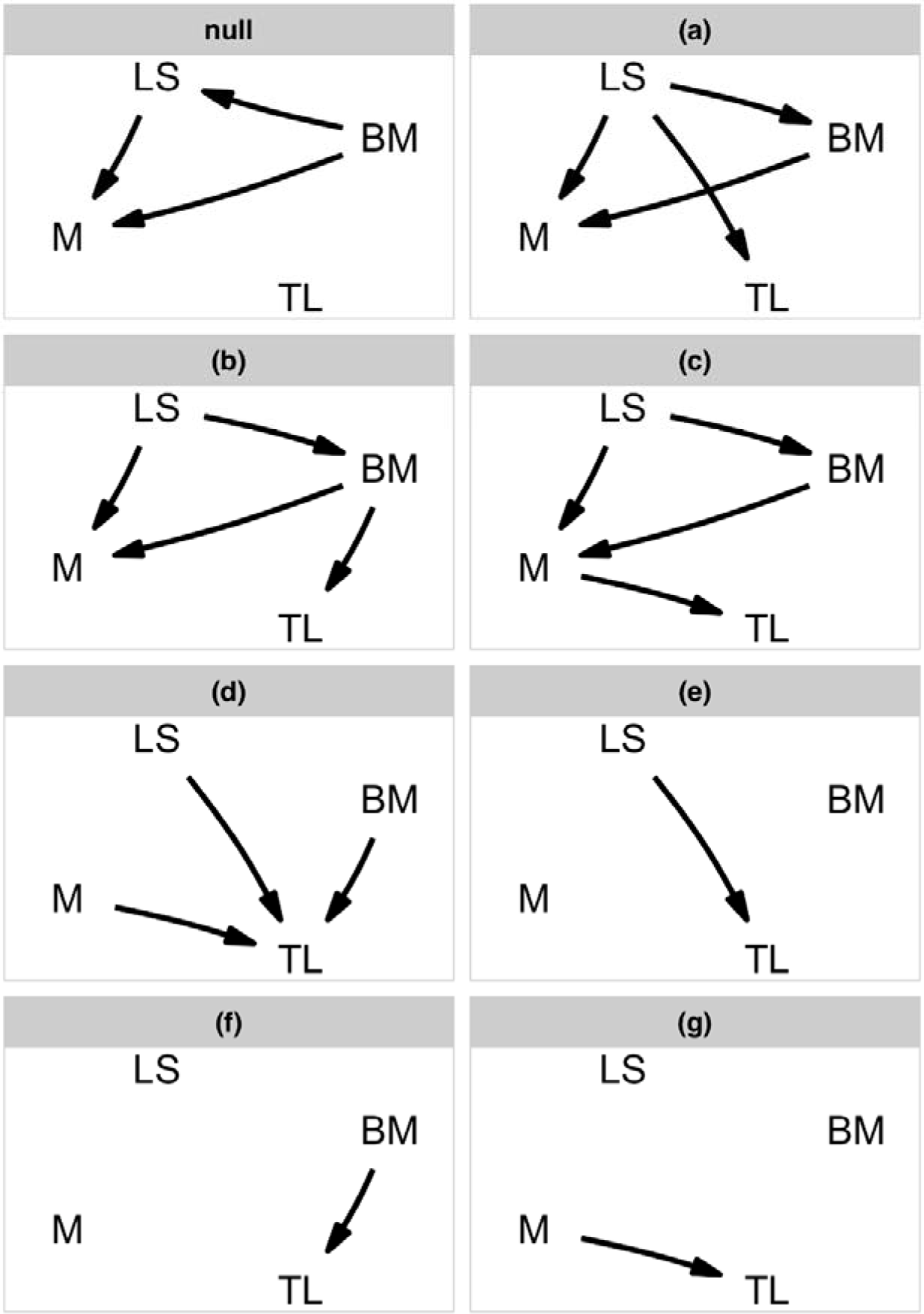
Path models depicting relationships among telomere length and predictor variables derived from optimality models (see text for details). Abbreviation in the path diagrams are as follows: TL, telomere length; BM, body mass; LS, lifespan; M, baseline metabolism.

**Figure S4:**
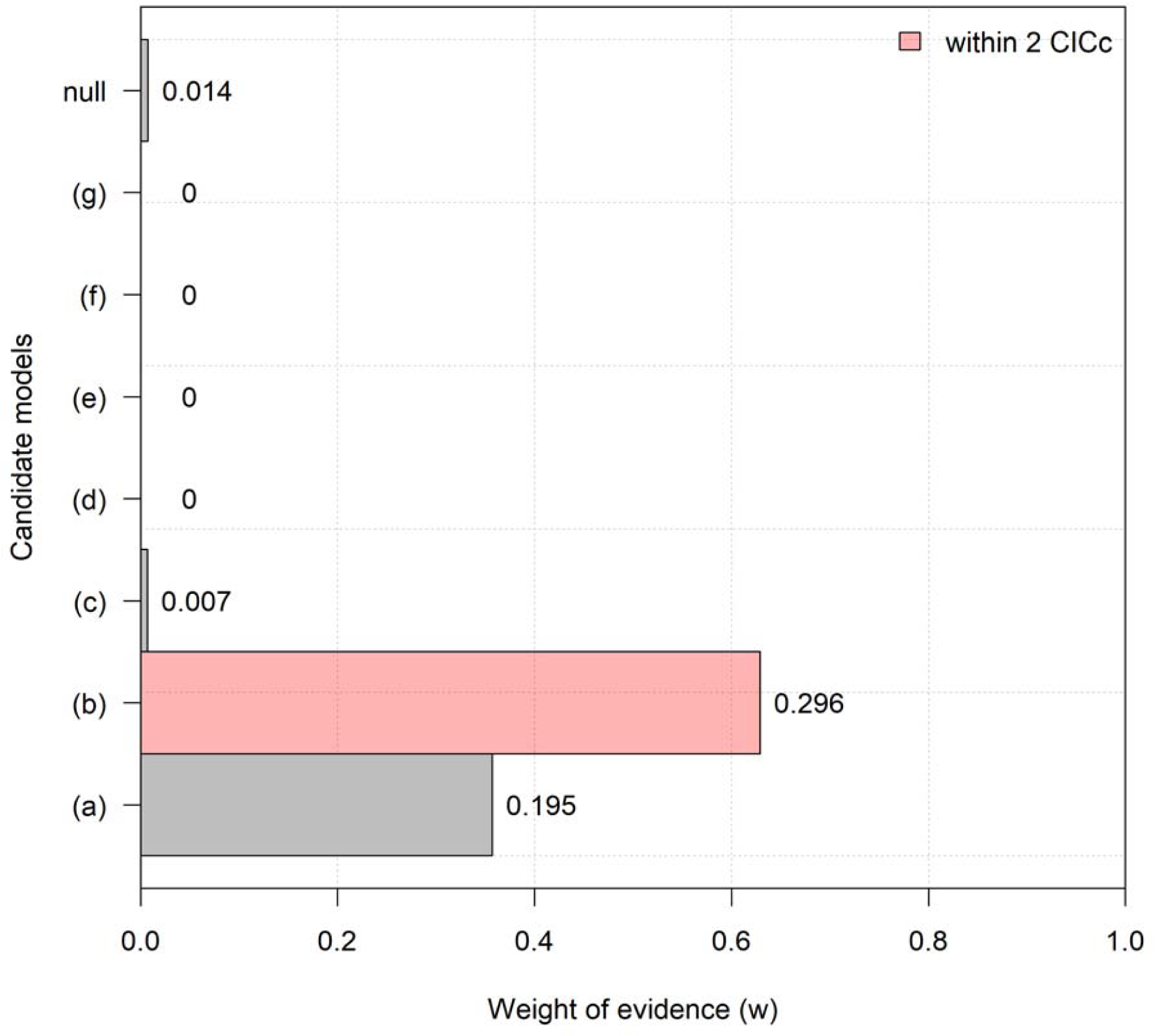
Relative importance of the causal models describing the evolution of telomere length across vertebrates. Bar labels are p-values; significance indicates rejection.

**Figure S5:**
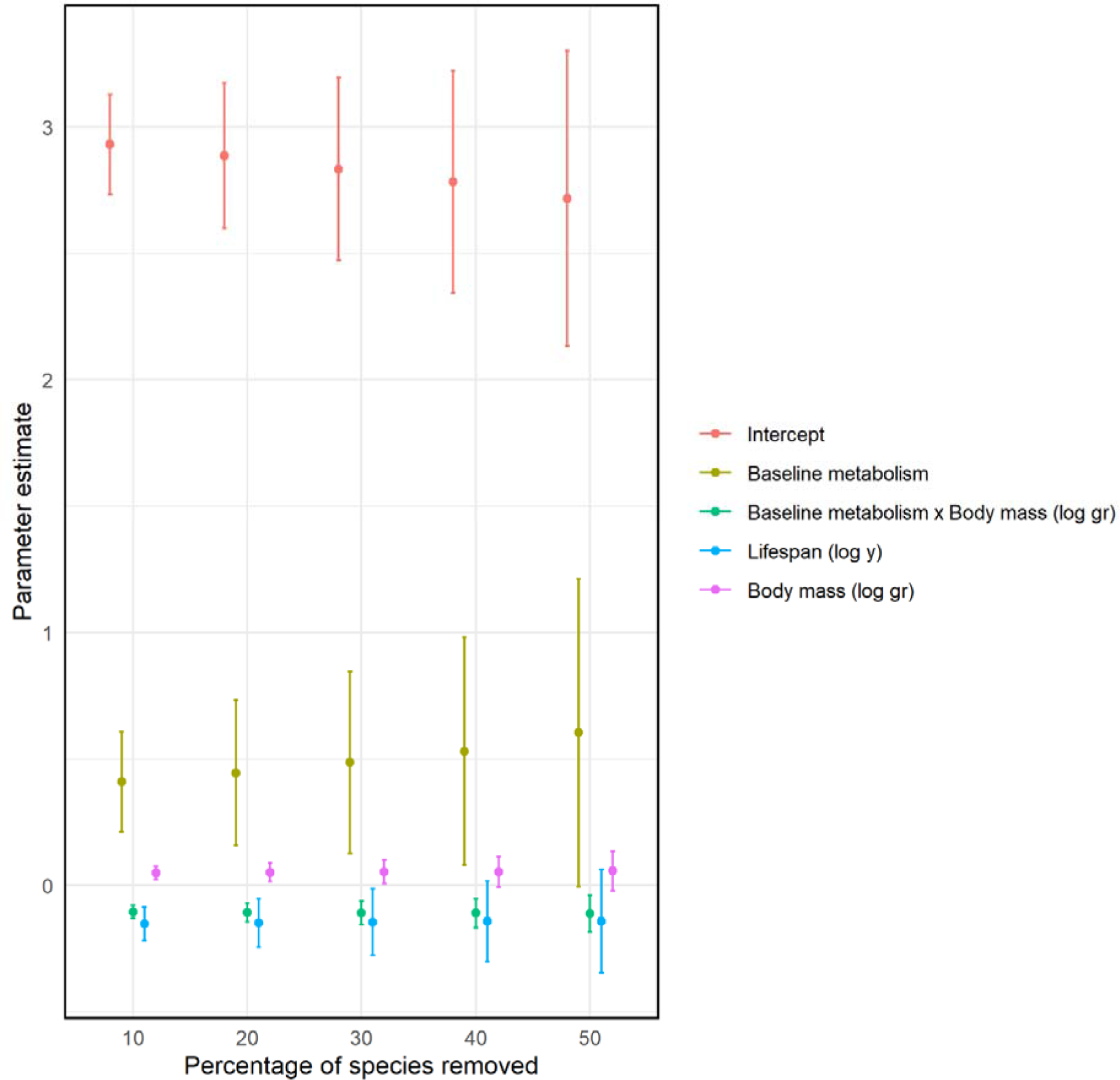
Sensitivity analysis investigating the effect of sample size on the parameter estimate of the intercept (red), body mass (purple), lifespan (blue), baseline metabolism (tan), and the baseline metabolism body mass interaction (green) on telomere length. Error bars represent the 95% CI of each parameter. 95% CIs not including zero indicates significance.

## References

Ahmed, W., & Lingner, J. (2018). Impact of oxidative stress on telomere biology. Differentiation, 99, 21–27. 10.1016/j.diff.2017.12.002

Apfelbeck, B., Haussmann, M. F., Boner, W., Flinks, H., Griffiths, K., Illera, J. C., Mortega, K. G., Sisson, Z., Smiddy, P., & Helm, B. (2019). Divergent patterns of telomere shortening in tropical compared to temperate stonechats. Ecology and Evolution, 9(1), 511–521. 10.1002/ece3.4769

Armstrong, E., & Boonekamp, J. (2023). Does oxidative stress shorten telomeres *in vivo*? A meta-analysis. Ageing Research Reviews, 85, 101854. 10.1016/j.arr.2023.101854

Aviv, A., Anderson, J. J., & Shay, J. W. (2017). Mutations, cancer and the telomere length paradox. Trends in Cancer, 3(4), 253–258. 10.1016/j.trecan.2017.02.005

Aubert, G., & Lansdorp, P. M. (2008). Telomeres and aging. Physiological Reviews, 88(2), 557–579. 10.1152/physrev.00026.2007

Baker, J., Meade, A., Pagel, M., & Venditti, C. (2021). Nothing wrong with the analysis of clades in comparative evolutionary studies: A reply to Poe et al. Systematic Biology, 70(1), 197–201. 10.1093/sysbio/syaa067

Bijl, W. van der. (2018). phylopath: Easy phylogenetic path analysis in R. PeerJ, 6, e4718. 10.7717/peerj.4718

Bonetti, D., Martina, M., Falcettoni, M., & Longhese, M. P. (2014). Telomere-end processing: Mechanisms and regulation. Chromosoma, 123(1), 57–66. 10.1007/s00412-013-0440-y

Cong, Y.-S., Wright, W. E., & Shay, J. W. (2002). Human telomerase and its regulation. Microbiology and Molecular Biology Reviews, 66(3), 407–425. 10.1128/mmbr.66.3.407-425.2002

Criscuolo, F., Dobson, F. S., & Schull, Q. (2021). The influence of phylogeny and life history on telomere lengths and telomere rate of change among bird species: A meta-analysis. Ecology and Evolution, 11(19), 12908–12922. 10.1002/ece3.7931

de Magalhães, J. P., Abidi, Z., dos Santos, G. A., Avelar, R. A., Barardo, D., Chatsirisupachai, K., Clark, P., De-Souza, E. A., Johnson, E. J., Lopes, I., Novoa, G., Senez, L., Talay, A., Thornton, D., & To, P. K. P. (2024). Human Ageing Genomic Resources: Updates on key databases in ageing research. Nucleic Acids Research, 52(D1), D900–D908. 10.1093/nar/gkad927

Elmore, L. W., Norris, M. W., Sircar, S., Bright, A. T., McChesney, P. A., Winn, R. N., & Holt, S. E. (2008). Upregulation of telomerase function during tissue regeneration. *Experimental Biology and Medicine (Maywood*, N.J.), 233(8), 958–967. 10.3181/0712-RM-345

Felsenstein, J. (2012). A comparative method for both discrete and continuous characters using the threshold model. The American Naturalist, 179(2), 145–156. 10.1086/663681

Fohringer, C., Hoelzl, F., Allen, A. M., Cayol, C., Ericsson, G., Spong, G., Smith, S., & Singh, N. J. (2022). Large mammal telomere length variation across ecoregions. BMC Ecology and Evolution, 22(1), 105. 10.1186/s12862-022-02050-5

Gilson, E., & Géli, V. (2007). How telomeres are replicated. Nature Reviews Molecular Cell Biology, 8(10), Article 10. 10.1038/nrm2259

Gokarn, R., Solon-Biet, S., Youngson, N. A., Wahl, D., Cogger, V. C., McMahon, A. C., Cooney, G. J., Ballard, J. W. O., Raubenheimer, D., Morris, M. J., Simpson, S. J., & Le Couteur, D. G. (2018). The relationship between dietary macronutrients and hepatic telomere length in aging mice. The Journals of Gerontology: Series A, 73(4), 446–449. 10.1093/gerona/glx186

Gomes, N. M. V., Ryder, O. A., Houck, M. L., Charter, S. J., Walker, W., Forsyth, N. R., Austad, S. N., Venditti, C., Pagel, M., Shay, J. W., & Wright, W. E. (2011). Comparative biology of mammalian telomeres: Hypotheses on ancestral states and the roles of telomeres in longevity determination. Aging Cell, 10(5), 761–768. 10.1111/j.1474-9726.2011.00718.x

Gomes, N. M. V., Shay, J. W., & Wright, W. E. (2010). Telomere biology in Metazoa. FEBS Letters, 584(17), 3741–3751. 10.1016/j.febslet.2010.07.031

Haussmann, M. F., & Marchetto, N. M. (2010). Telomeres: Linking stress and survival, ecology and evolution. Current Zoology, 56(6), 714–727. 10.1093/czoolo/56.6.714

Haussmann, M. F., Winkler, D. W., O’Reilly, K. M., Huntington, C. E., Nisbet, I. C. T., & Vleck, C. M. (2003). Telomeres shorten more slowly in long-lived birds and mammals than in short–lived ones. Proceedings of the Royal Society of London. Series B: Biological Sciences, 270(1522), 1387–1392. 10.1098/rspb.2003.2385

Hayes, J. P. (2010). Metabolic rates, genetic constraints, and the evolution of endothermy. Journal of Evolutionary Biology, 23(9), 1868–1877. 10.1111/j.1420-9101.2010.02053.x

Heidinger, B. J., Blount, J. D., Boner, W., Griffiths, K., Metcalfe, N. B., & Monaghan, P. (2012). Telomere length in early life predicts lifespan. Proceedings of the National Academy of Sciences, 109(5), 1743–1748. 10.1073/pnas.1113306109

Jimenez, A. G., Harper, J. M., Queenborough, S. A., & Williams, J. B. (2013). Linkages between the life-history evolution of tropical and temperate birds and the resistance of cultured skin fibroblasts to oxidative and non-oxidative chemical injury. Journal of Experimental Biology, 216(8), 1373–1380. 10.1242/jeb.079889

Kawanishi, S., & Oikawa, S. (2004). Mechanism of telomere shortening by oxidative stress. Annals of the New York Academy of Sciences, 1019(1), 278–284. 10.1196/annals.1297.047

Lange, T. de. (2005). Shelterin: The protein complex that shapes and safeguards human telomeres. Genes & Development, 19(18), 2100–2110. 10.1101/gad.1346005

McLester-Davis, L. W. Y., Estrada, P., Hastings, W. J., Kataria, L. A., Martin, N. A., Siebeneicher, J. T., Tristano, R. I., Mayne, C. V., Horlick, R. P., O’Connell, S. S., & Drury, S. S. (2023). A review and meta-analysis: Cross-tissue telomere length correlations in healthy humans. Ageing Research Reviews, 88, 101942. 10.1016/j.arr.2023.101942

McNally, E. J., Luncsford, P. J., & Armanios, M. (2019). Long telomeres and cancer risk: The price of cellular immortality. The Journal of Clinical Investigation, 129(9), 3474–3481. 10.1172/JCI120851

Moretton, A., & Loizou, J. I. (2020). Interplay between Cellular Metabolism and the DNA Damage Response in Cancer. Cancers, 12(8), Article 8. 10.3390/cancers12082051

Neumann, A. A., Watson, C. M., Noble, J. R., Pickett, H. A., Tam, P. P. L., & Reddel, R. R. (2013). Alternative lengthening of telomeres in normal mammalian somatic cells. Genes & Development, 27(1), 18–23. 10.1101/gad.205062.112

Olsson, M., Wapstra, E., & Friesen, C. (2018). Ectothermic telomeres: It’s time they came in from the cold. Philosophical Transactions of the Royal Society B: Biological Sciences, 373(1741), 20160449. 10.1098/rstb.2016.0449

Paul, L. (2011). Diet, nutrition and telomere length. The Journal of Nutritional Biochemistry, 22(10), 895–901. 10.1016/j.jnutbio.2010.12.001

Pepke, M. L., & Eisenberg, D. T. A. (2022). On the comparative biology of mammalian telomeres: Telomere length co-evolves with body mass, lifespan and cancer risk. Molecular Ecology, 31(23), 6286–6296. 10.1111/mec.15870

Pepke, M. L., Kvalnes, T., Wright, J., Araya-Ajoy, Y. G., Ranke, P. S., Boner, W., Monaghan, P., Sæther, B.-E., Jensen, H., & Ringsby, T. H. (2023b). Longitudinal telomere dynamics within natural lifespans of a wild bird. Scientific Reports, 13(1), Article 1. 10.1038/s41598-023-31435-9

Pepke, M. L., Ringsby, T. H., & Eisenberg, D. T. A. (2023a). The evolution of early-life telomere length, pace-of-life and telomere-chromosome length dynamics in birds. Molecular Ecology, 32(11), 2898–2912. 10.1111/mec.16907

Peto, R., Roe, F. J., Lee, P. N., Levy, L., & Clack, J. (1975). Cancer and ageing in mice and men. British Journal of Cancer, 32(4), Article 4. 10.1038/bjc.1975.242

R Core Team (2022). R: A language and environment for statistical computing. R Foundation for Statistical Computing. https://www.R-project.org

Revell, L. J., & Harmon, L. J. (2022). Phylogenetic comparative methods in R. Princeton University Press.

Rhodes, D., & Giraldo, R. (1995). Telomere structure and function. Current Opinion in Structural Biology, 5(3), 311–322. 10.1016/0959-440X(95)80092-1

Risques, R. A., & Promislow, D. E. L. (2018). All’s well that ends well: Why large species have short telomeres. Philosophical Transactions of the Royal Society B: Biological Sciences, 373(1741), 20160448. 10.1098/rstb.2016.0448

Rocco, L., Costagliola, D., & Stingo, V. (2001). (TTAGGG)n telomeric sequence in selachian chromosomes. Heredity, 87(5), Article 5. 10.1046/j.1365-2540.2001.00945.x

Rode, L., Nordestgaard, B. G., & Bojesen, S. E. (2016). Long telomeres and cancer risk among 95LJ568 individuals from the general population. International Journal of Epidemiology, 45(5), 1634–1643. 10.1093/ije/dyw179

Rozhok, A. I., & DeGregori, J. (2016). The evolution of lifespan and age-dependent cancer risk. Trends in Cancer, 2(10), 552–560. 10.1016/j.trecan.2016.09.004

Saretzki, G., & Von Zglinicki, T. (2002). Replicative aging, telomeres, and oxidative stress. Annals of the New York Academy of Sciences, 959(1), 24–29. 10.1111/j.1749-6632.2002.tb02079.x

Schneeberger, K., Czirják, G. Á., & Voigt, C. C. (2014). Frugivory is associated with low measures of plasma oxidative stress and high antioxidant concentration in free-ranging bats. Die Naturwissenschaften, 101(4), 285–290. 10.1007/s00114-014-1155-5

Seluanov, A., Chen, Z., Hine, C., Sasahara, T. H. C., Ribeiro, A. A. C. M., Catania, K. C., Presgraves, D. C., & Gorbunova, V. (2007). Telomerase activity coevolves with body mass not lifespan. Aging Cell, 6(1), 45–52. 10.1111/j.1474-9726.2006.00262.x

Seluanov, A., Hine, C., Bozzella, M., Hall, A., Sasahara, T. H. C., Ribeiro, A. A. C. M., Catania, K. C., Presgraves, D. C., & Gorbunova, V. (2008). Distinct tumor suppressor mechanisms evolve in rodent species that differ in size and lifespan. Aging Cell, 7(6), 813–823. 10.1111/j.1474-9726.2008.00431.x

Shay, J. W., & S. A. Bacchetti. (1997). A survey of telomerase activity in human cancer. European Journal of Cancer, 33, 787–791. 10.1016/S0959-8049(97)00062-2

Shay, J. W., & Wright, W. E. (2005). Senescence and immortalization: Role of telomeres and telomerase. Carcinogenesis, 26(5), 867–874. 10.1093/carcin/bgh296

Shay, J. W., & Wright, W. E. (2011). Role of telomeres and telomerase in cancer. Seminars in Cancer Biology, 21(6), 349–353. 10.1016/j.semcancer.2011.10.001

Shipley, B. (2013). The AIC model selection method applied to path analytic models compared using a d-separation test. Ecology, 94(3), 560–564. 10.1890/12-0976.1

Stone, E. A. (2011). Why the phylogenetic regression appears robust to tree misspecification. Systematic Biology, 60(3), 245–260. 10.1093/sysbio/syq098

Sun, J., Liu, W., Guo, Y., Zhang, H., Jiang, D., Luo, Y., Liu, R., & Chen, C. (2021). Characterization of tree shrew telomeres and telomerase. Journal of Genetics and Genomics, 48(7), 631–639. 10.1016/j.jgg.2021.06.004

Tricola, G. M., Simons, M. J. P., Atema, E., Boughton, R. K., Brown, J. L., Dearborn, D. C., Divoky, G., Eimes, J. A., Huntington, C. E., Kitaysky, A. S., Juola, F. A., Lank, D. B., Litwa, H. P., Mulder, E. G. A., Nisbet, I. C. T., Okanoya, K., Safran, R. J., Schoech, S. J., Schreiber, E. A., … Haussmann, M. F. (2018). The rate of telomere loss is related to maximum lifespan in birds. Philosophical Transactions of the Royal Society B: Biological Sciences, 373(1741), 20160445. 10.1098/rstb.2016.0445

Watson, H., Bolton, M., & Monaghan, P. (2015). Variation in early-life telomere dynamics in a long-lived bird: Links to environmental conditions and survival. Journal of Experimental Biology, 218(5), 668–674. 10.1242/jeb.104265

Whittemore, K., Vera, E., Martínez-Nevado, E., Sanpera, C., & Blasco, M. A. (2019). Telomere shortening rate predicts species life span. Proceedings of the National Academy of Sciences, 116(30), 15122–15127. 10.1073/pnas.1902452116

Wright, W. E., Piatyszek, M. A., Rainey, W. E., Byrd, W., & Shay, J. W. (1996). Telomerase activity in human germline and embryonic tissues and cells. Developmental Genetics, 18(2), 173–179. 10.1002/(SICI)1520-6408(1996)18:2<173::AID-DVG10>3.0.CO;2-3

Xu, L., Li, S., & Stohr, B. A. (2013). The role of telomere biology in cancer. Annual Review of Pathology: Mechanisms of Disease, 8(1), 49–78. 10.1146/annurev-pathol-020712-164030

Zglinicki, T. von. (2002). Oxidative stress shortens telomeres. Trends in Biochemical Sciences, 27(7), 339–344. 10.1016/S0968-0004(02)02110-2

Zhdanova, N. S., Karamisheva, T. V., Minina, J., Astakhova, N. M., Lansdorp, P., Kammori, M., Rubtsov, N. B., & Searle, J. B. (2005). Unusual distribution pattern of telomeric repeats in the shrews *Sorex araneus* and *Sorex granarius*. Chromosome Research, 13(6), 617–625. 10.1007/s10577-005-0988-3

