## Supplementary table 1 references for "Why are telomeres the length that they are? Insight from a phylogenetic comparative analysis"

Allen, G. R. (1989). Freshwater Fishes of Australia TFH Publications. *Inc., Neptune City, New Jersey*.

Anchelin, M., Murcia, L., Alcaraz-Pérez, F., García-Navarro, E. M., & Cayuela, M. L. (2011). Behaviour of telomere and telomerase during aging and regeneration in Zebrafish. *PLOS ONE*, *6*(2), e16955. <https://doi.org/10.1371/journal.pone.0016955>

Bauch, C., Gatt, M. C., Granadeiro, J. P., Verhulst, S., & Catry, P. (2020). Sex‐specific telomere length and dynamics in relation to age and reproductive success in Cory’s shearwaters. *Molecular Ecology*, *29*(7), 1344–1357. <https://doi.org/10.1111/mec.15399>

Belmaker, A. (2016). The Role Of Telomere Length In Tree Swallow Behavior And Life History. <https://doi.org/10.7298/X4J38QG8>

Belmaker, A., Hallinger, K. K., Glynn, R. A., Winkler, D. W., & Haussmann, M. F. (2019). The environmental and genetic determinants of chick telomere length in tree swallows (*Tachycineta bicolor*). *Ecology and Evolution*, *9*(14), 8175–8186. <https://doi.org/10.1002/ece3.5386>

Berman, D. I., Bulakhova, N. A., Alfimov, A. V., & Meshcheryakova, E. N. (2016). How the most northern lizard, Zootoca vivipara, overwinters in Siberia. *Polar Biology*, *39*(12), 2411–2425. <https://doi.org/10.1007/s00300-016-1916-z>

Bhatnagar, Y. M., Mishra, P. V., Martinez, J. A., Saxena, K. M., & Wertelecki, W. (1995). Telomeres of higher primates. *Biochemistry and Molecular Biology International*, *37*(1), 57–64.

Bichet, C., Bouwhuis, S., Bauch, C., Verhulst, S., Becker, P. H., & Vedder, O. (2020). Telomere length is repeatable, shortens with age and reproductive success, and predicts remaining lifespan in a long-lived seabird. *Molecular Ecology*, *29*(2), 429–441. <https://doi.org/10.1111/mec.15331>

Berman, D. I., Bulakhova, N. A., Alfimov, A. V., & Meshcheryakova, E. N. (2016). How the most northern lizard, *Zootoca vivipara*, overwinters in Siberia. *Polar Biology*, *39*(12), 2411–2425. <https://doi.org/10.1007/s00300-016-1916-z>

Blévin, P., Angelier, F., Tartu, S., Bustamante, P., Herzke, D., Moe, B., Bech, C., Gabrielsen, G. W., Bustnes, J. O., & Chastel, O. (2017). Perfluorinated substances and telomeres in an Arctic seabird: Cross-sectional and longitudinal approaches. *Environmental Pollution*, *230*, 360–367. <https://doi.org/10.1016/j.envpol.2017.06.060>

Caprioli, M., Romano, M., Romano, A., Rubolini, D., Motta, R., Folini, M., & Saino, N. (2013). Nestling telomere length does not predict longevity, but covaries with adult body size in wild barn swallows. *Biology Letters*, *9*(5), 20130340. <https://doi.org/10.1098/rsbl.2013.0340>

Chochi, M., Niizuma, Y., & Takagi, M. (2002). Sexual differences in the external measurements of black-tailed gulls breeding on Rishiri Island, Japan. *Ornithological Science*, *1*(2), 163–166. <https://doi.org/10.2326/osj.1.163>

Clark, T. S., Pandolfo, L. M., Marshall, C. M., Mitra, A. K., & Schech, J. M. (2018). Body Condition Scoring for Adult Zebrafish (Danio rerio). *Journal of the American Association for Laboratory Animal Science*, *57*(6), 698–702. <https://doi.org/10.30802/AALAS-JAALAS-18-000045>

Criscuolo, F., Bize, P., Nasir, L., Metcalfe, N. B., Foote, C. G., Griffiths, K., Gault, E. A., & Monaghan, P. (2009). Real-time quantitative PCR assay for measurement of avian telomeres. *Journal of Avian Biology*, *40*(3), 342–347. <https://doi.org/10.1111/j.1600-048X.2008.04623.x>

Criscuolo, F., Dobson, F. S., & Schull, Q. (2021). The influence of phylogeny and life history on telomere lengths and telomere rate of change among bird species: A meta-analysis. *Ecology and Evolution*, *11*(19), 12908–12922. <https://doi.org/10.1002/ece3.7931>

de Abechuco, E. L., Bilbao, E., Soto, M., & Díez, G. (2014). Molecular cloning and measurement of telomerase reverse transcriptase (TERT) transcription patterns in tissues of European hake (*Merluccius merluccius*) and Atlantic cod (*Gadus morhua*) during aging. *Gene*, *541*(1), 8–18. <https://doi.org/10.1016/j.gene.2014.03.006>

de Abechuco, E. L., Hartmann, N., Soto, M., & Díez, G. (2016). Assessing the variability of telomere length measures by means of Telomeric Restriction Fragments (TRF) in different tissues of cod *Gadus morhua*. *Gene Reports*, *5*, 117–125. <https://doi.org/10.1016/j.genrep.2016.09.009>

de Magalhães, J. P., Abidi, Z., dos Santos, G. A., Avelar, R. A., Barardo, D., Chatsirisupachai, K., Clark, P., De-Souza, E. A., Johnson, E. J., Lopes, I., Novoa, G., Senez, L., Talay, A., Thornton, D., & To, P. K. P. (2024). Human Ageing Genomic Resources: Updates on key databases in ageing research. *Nucleic Acids Research*, *52*(D1), D900–D908. <https://doi.org/10.1093/nar/gkad927>

de Magalhães, J. P., Abidi, Z., dos Santos, G. A., Avelar, R. A., Barardo, D., Chatsirisupachai, K., Clark, P., De-Souza, E. A., Johnson, E. J., Lopes, I., Novoa, G., Senez, L., Talay, A., Thornton, D., & To, P. K. P. (2024). Human Ageing Genomic Resources: Updates on key databases in ageing research. *Nucleic Acids Research*, *52*(D1), D900–D908. <https://doi.org/10.1093/nar/gkad927>

Delany, M. E., Krupkin, A. B., & Miller, M. M. (2000). Organization of telomere sequences in birds: Evidence for arrays of extreme length and for in vivo shortening. *Cytogenetics and Cell Genetics*, *90*(1–2), 139–145. <https://doi.org/10.1159/000015649>

Downs, K. P., Shen, Y., Pasquali, A., Beldorth, I., Savage, M., Gallier, K., Garcia, T., Booth, R. E., & Walter, R. B. (2012). Characterization of telomeres and telomerase expression in *Xiphophorus*. *Comparative Biochemistry and Physiology Part C: Toxicology & Pharmacology*, *155*(1), 89–94. <https://doi.org/10.1016/j.cbpc.2011.05.005>

Dupoué, A., Rutschmann, A., Le Galliard, J. F., Clobert, J., Angelier, F., Marciau, C., Ruault, S., Miles, D., & Meylan, S. (2017). Shorter telomeres precede population extinction in wild lizards. *Scientific Reports*, *7*(1), Article 1. <https://doi.org/10.1038/s41598-017-17323-z>

Escoriza, D., Poch, S., Sunyer-Sala, P., & Boix, D. (2021). Growth patterns of *Emys orbicularis* across a range of aquatic habitats: A long-term study. *Basic and Applied Herpetology*, *35*, 0–000. <https://doi.org/10.11160/bah.228>

Feng, Y.-R., Norwood Jr., D., Shibata, R., Gee, D., Xiao, X., Martin, M., Zeichner, S. L., & Dimitrov, D. S. (1998). Telomere dynamics in HIV-1 infected and uninfected chimpanzees measured by an improved method based on high-resolution two-dimensional calibration of DNA sizes. *Journal of Medical Primatology*, *27*(5), 258–265. <https://doi.org/10.1111/j.1600-0684.1998.tb00246.x>

*FishBase: A Global Information System on Fishes*. (n.d.). <https://fishbase.se/home.htm>

Fongy, A., Romestaing, C., Blanc, C., Lacoste-Garanger, N., Rouanet, J.-L., Raccurt, M., & Duchamp, C. (2013). Ontogeny of muscle bioenergetics in Adélie penguin chicks (*Pygoscelis adeliae*). *American Journal of Physiology-Regulatory, Integrative and Comparative Physiology*, *305*(9), R1065–R1075. <https://doi.org/10.1152/ajpregu.00137.2013>

Foote, C. (2009). *Avian telomere dynamics*. <https://www.semanticscholar.org/paper/Avian-telomere-dynamics-Foote/9eae45af2677491a47d98ed222592e4e7efa6051>

Foote, C. G., Daunt, F., González-Solís, J., Nasir, L., Phillips, R. A., & Monaghan, P. (2010). Individual state and survival prospects: Age, sex, and telomere length in a long-lived seabird. *Behavioral Ecology*, *22*(1), 156–161. <https://doi.org/10.1093/beheco/arq178>

Foote, C. G., Gault, E. A., Nasir, L., & Monaghan, P. (2011). Telomere dynamics in relation to early growth conditions in the wild in the lesser black-backed gull. *Journal of Zoology*, *283*(3), 203–209. <https://doi.org/10.1111/j.1469-7998.2010.00774.x>

Foote, C. G., Vleck, D., & Vleck, C. M. (2013). Extent and variability of interstitial telomeric sequences and their effects on estimates of telomere length. *Molecular Ecology Resources*, *13*(3), 417–428. <https://doi.org/10.1111/1755-0998.12079>

Forsyth, N. R., Elder, F. F. B., Shay, J. W., & Wright, W. E. (2005). Lagomorphs (rabbits, pikas and hares) do not use telomere-directed replicative aging in vitro. *Mechanisms of Ageing and Development*, *126*(6), 685–691. <https://doi.org/10.1016/j.mad.2005.01.003>

Gibbons, J. W., & Semlitsch, R. D. (1982). Survivorship and longevity of a long-lived vertebrate species: How long do turtles live? *Journal of Animal Ecology*, *51*(2), 523–527. <https://doi.org/10.2307/3981>

Girondot, M. and Garcia, J. (1999) Senescence and longevity in turtles: what telomeres tell us. in Proceedings of the 9th Ordinary General Meeting of the Societas Europaea Herpetologica In Current Studies in Herpetology, ed Guyetant CMaR (Le Bourget du Lac, France), pp 133–137.

Gomes, N. M. V., Ryder, O. A., Houck, M. L., Charter, S. J., Walker, W., Forsyth, N. R., Austad, S. N., Venditti, C., Pagel, M., Shay, J. W., & Wright, W. E. (2011). Comparative biology of mammalian telomeres: Hypotheses on ancestral states and the roles of telomeres in longevity determination. *Aging Cell*, *10*(5), 761–768. <https://doi.org/10.1111/j.1474-9726.2011.00718.x>

Guo, Y.-Z., Zhang, Y., Wang, Q., Yu, J., Wan, Q.-H., Huang, J., & Fang, S.-G. (2023). Alternative telomere maintenance mechanism in Alligator sinensis provides insights into aging evolution. *iScience*, *26*(1), 105850. <https://doi.org/10.1016/j.isci.2022.105850>

Hall, M. E., Nasir, L., Daunt, F., Gault, E. A., Croxall, J. P., Wanless, S., & Monaghan, P. (2004). Telomere loss in relation to age and early environment in long-lived birds. *Proceedings of the Royal Society of London. Series B: Biological Sciences*, *271*(1548), 1571–1576. <https://doi.org/10.1098/rspb.2004.2768>

Hartmann, N., Reichwald, K., Lechel, A., Graf, M., Kirschner, J., Dorn, A., Terzibasi, E., Wellner, J., Platzer, M., Rudolph, K. L., Cellerino, A., & Englert, C. (2009). Telomeres shorten while Tert expression increases during ageing of the short-lived fish *Nothobranchius furzeri*. *Mechanisms of Ageing and Development*, *130*(5), 290–296. <https://doi.org/10.1016/j.mad.2009.01.003>

Haussmann, M. F., Winkler, D. W., O’Reilly, K. M., Huntington, C. E., Nisbet, I. C. T., & Vleck, C. M. (2003). Telomeres shorten more slowly in long-lived birds and mammals than in short–lived ones. *Proceedings of the Royal Society of London. Series B: Biological Sciences*, *270*(1522), 1387–1392. <https://doi.org/10.1098/rspb.2003.2385>

Haussmann, M. F., Winkler, D. W., Huntington, C. E., Nisbet, I. C. T., & Vleck, C. M. (2004). Telomerase expression is differentially regulated in birds of differing life span. *Annals of the New York Academy of Sciences*, *1019*(1), 186–190. <https://doi.org/10.1196/annals.1297.029>

Haussmann, M. F., Winkler, D. W., Huntington, C. E., Nisbet, I. C., & Vleck, C. M. (2007). Telomerase activity is maintained throughout the lifespan of long-lived birds. *Experimental Gerontology*, *42*(7), 610–618.

Horn, T., Gemmell, N. J., Robertson, B. C., & Bridges, C. R. (2008). Telomere length change in European sea bass (*Dicentrarchus labrax*). *Australian Journal of Zoology*, *56*(3), 207–210. <https://doi.org/10.1071/ZO08046>

Horn, T., Robertson, B. C., Will, M., Eason, D. K., Elliott, G. P., & Gemmell, N. J. (2011). Inheritance of telomere length in a bird. *PLOS ONE*, *6*(2), e17199. <https://doi.org/10.1371/journal.pone.0017199>

Hsu, C.-Y., Chiu, Y.-C., Hsu, W.-L., & Chan, Y.-P. (2008). Age-related markers assayed at different developmental stages of the annual fish *Nothobranchius rachovii*. *The Journals of Gerontology: Series A*, *63*(12), 1267–1276. <https://doi.org/10.1093/gerona/63.12.1267>

Hyodo, S., Bell, J. D., Healy, J. M., Kaneko, T., Hasegawa, S., Takei, Y., Donald, J. A., & Toop, T. (2007). Osmoregulation in elephant fish *Callorhinchus milii* (Holocephali), with special reference to the rectal gland. *Journal of Experimental Biology*, *210*(8), 1303–1310. <https://doi.org/10.1242/jeb.003418>

Ibáñez-Álamo, J. D., Pineda-Pampliega, J., Thomson, R. L., Aguirre, J. I., Díez-Fernández, A., Faivre, B., Figuerola, J., & Verhulst, S. (2018). Urban blackbirds have shorter telomeres. *Biology Letters*, *14*(3), 20180083. <https://doi.org/10.1098/rsbl.2018.0083>

IGFA, 2001. Database of IGFA angling records until 2001. IGFA, Fort Lauderdale, USA.

Izzo, C. (2010). Patterns of telomere length change with age in aquatic vertebrates and the phylogenetic distribution of the pattern among jawed vertebrates*.* [Thesis]. <https://digital.library.adelaide.edu.au/dspace/handle/2440/63477>

Kailola, P. J., Williams, M. J., Stewart, P. C., Reichelt, R. E., & McNee, A. (1993). Australian fisheries resources.

Koopman, Matt & Morison, Sandy & Victoria. Department of Primary Industries & Victoria. Marine and Freshwater Resources Institute. (2002). Sand flathead - a bread and butter catch [electronic resource] / Matt Koopman and Sandy Morison, Marine and Freshwater Resources Institute. [Melbourne] : Dept. of Primary Industries

Law, C. S. W., & Sadovy de Mitcheson, Y. (2018). Age and growth of black seabream Acanthopagrus schlegelii (Sparidae) in Hong Kong and adjacent waters of the northern South China Sea. *Journal of Fish Biology*, *93*(2), 382–390. <https://doi.org/10.1111/jfb.13774>

Lejnine, S., Makarov, V. L., & Langmore, J. P. (1995). Conserved nucleoprotein structure at the ends of vertebrate and invertebrate chromosomes. *Proceedings of the National Academy of Sciences of the United States of America*, *92*(6), 2393–2397. <https://doi.org/10.1073/pnas.92.6.2393>

Marinewise. (n.d.). Australian marine life, boating, fishing & more. Marinewise. <https://marinewise.com.au/>

McKenzie, W. D., Crews, D., Kallman, K. D., Policansky, D., & Sohn, J. J. (1983). Age, weight and the genetics of sexual maturation in the platyfish, *Xiphophorus maculatus*. *Copeia*, *1983*(3), 770–774. <https://doi.org/10.2307/1444344>

Meyer, C. I., Kaufman, R., & Cech, J. J. (2006). Melanin pattern morphs do not differ in metabolic rate: Implications for the evolutionary maintenance of a melanophore polymorphism in the green swordtail, Xiphophorus helleri. *Naturwissenschaften*, *93*(10), 495–499. <https://doi.org/10.1007/s00114-006-0134-x>

Mizutani, Y., Tomita, N., Niizuma, Y., & Yoda, K. (2013). Environmental perturbations influence telomere dynamics in long-lived birds in their natural habitat. *Biology Letters*, *9*(5), 20130511. <https://doi.org/10.1098/rsbl.2013.0511>

Moore, O. R., Stutchbury, B. J. M., & Quinn, J. S. (1999). Extrapair mating system of an asynchronously breeding tropical songbird: The mangrove swallow. *The Auk*, *116*(4), 1039–1049. <https://doi.org/10.2307/4089683>

*Museum of Zoology | U-M LSA Museum of Zoology*. (n.d.). <https://lsa.umich.edu/ummz>

Nowak, Ronald M., and J. L. Paradiso. "Mammals of the World." *John Hopkins Press, Baltimore* (1991).

Olsson, M., Gullberg, A., & Tegelström, H. (1997). Determinants of breeding dispersal in the sand lizard, Lacerta agilis, (Reptilia, Squamata). *Biological Journal of the Linnean Society*, *60*(2), 243–256. <https://doi.org/10.1111/j.1095-8312.1997.tb01494.x>

Olsson, M., Pauliny, A., Wapstra, E., & Blomqvist, D. (2010). Proximate determinants of telomere length in sand lizards (*Lacerta agilis*). *Biology Letters*, *6*(5), 651–653. <https://doi.org/10.1098/rsbl.2010.0126>

Ouyang, J. Q., Lendvai, Á. Z., Moore, I. T., Bonier, F., & Haussmann, M. F. (2016). Do hormones, telomere lengths, and oxidative stress form an integrated phenotype? A case study in free-living tree swallows. *Integrative and Comparative Biology*, *56*(2), 138–145. <https://doi.org/10.1093/icb/icw044>

Panasiak, L., Kuciński, M., Hliwa, P., Pomianowski, K., & Ocalewicz, K. (2023). Telomerase Activity in Somatic Tissues and Ovaries of Diploid and Triploid Rainbow Trout (*Oncorhynchus mykiss*) Females. *Cells*, *12*(13), Article 13. <https://doi.org/10.3390/cells12131772>

Pauliny, A., Wagner, R. H., Augustin, J., Szép, T., & Blomqvist, D. (2006). Age-independent telomere length predicts fitness in two bird species. *Molecular Ecology*, *15*(6), 1681–1687. <https://doi.org/10.1111/j.1365-294X.2006.02862.x>

Pauliny, A., Larsson, K., & Blomqvist, D. (2012). Telomere dynamics in a long-lived bird, the barnacle goose. *BMC Evolutionary Biology*, *12*(1), 257. <https://doi.org/10.1186/1471-2148-12-257>

Polačik, M., & Reichard, M. (2010). Diet overlap among three sympatric African annual killifish species *Nothobranchius* spp. From Mozambique. *Journal of Fish Biology*, *77*(3), 754–768. <https://doi.org/10.1111/j.1095-8649.2010.02717.x>

Scott, N. M., Haussmann, M. F., Elsey, R. M., Trosclair, P. L., & Vleck, C. M. (2006). Telomere length shortens with body length in *Alligator mississippiensis*. *Southeastern Naturalist*, *5*(4), 685–692. [https://doi.org/10.1656/1528-7092(2006)5[685:TLSWBL]2.0.CO;2](https://doi.org/10.1656/1528-7092(2006)5%5b685:TLSWBL%5d2.0.CO;2)

Seluanov, A., Chen, Z., Hine, C., Sasahara, T. H. C., Ribeiro, A. A. C. M., Catania, K. C., Presgraves, D. C., & Gorbunova, V. (2007). Telomerase activity coevolves with body mass not lifespan. *Aging Cell*, *6*(1), 45–52. <https://doi.org/10.1111/j.1474-9726.2006.00262.x>

Shibata, R., Feng, Y.-R., Gee, D., Norwood, D., Xiao, X., Zeichner, S. L., Martin, M. A., & Dimitrov, D. S. (1999). Telomere dynamics in monkeys: Increased cell turnover in macaques infected with chimeric simian-human immunodeficiency viruses. *Journal of Medical Primatology*, *28*(1), 1–10. <https://doi.org/10.1111/j.1600-0684.1999.tb00083.x>

*Status of Australian Fish Stocks Reports*. (n.d.). Retrieved May 14, 2024, from <https://fish.gov.au/>

Steinert, S., White, D. M., Zou, Y., Shay, J. W., & Wright, W. E. (2002). Telomere Biology and Cellular Aging in Nonhuman Primate Cells. *Experimental Cell Research*, *272*(2), 146–152. <https://doi.org/10.1006/excr.2001.5409>

Tricola, G. M., Simons, M. J. P., Atema, E., Boughton, R. K., Brown, J. L., Dearborn, D. C., Divoky, G., Eimes, J. A., Huntington, C. E., Kitaysky, A. S., Juola, F. A., Lank, D. B., Litwa, H. P., Mulder, E. G. A., Nisbet, I. C. T., Okanoya, K., Safran, R. J., Schoech, S. J., Schreiber, E. A., … Haussmann, M. F. (2018). The rate of telomere loss is related to maximum lifespan in birds. *Philosophical Transactions of the Royal Society B: Biological Sciences*, *373*(1741), 20160445. <https://doi.org/10.1098/rstb.2016.0445>

Trinnie, F., Walker, T., Jones, P., & Laurenson, L. (2014). Regional differences in the reproductive parameters of the sparsely-spotted stingaree, *Urolophus paucimaculatus*, from South-Eastern Australia. *Marine and Freshwater Research*, *65*, 943–958. <https://doi.org/10.1071/MF13275>

Tsui, C. (2005). Evaluation of telomere length as an age-marker in marine teleosts. [*Http://Sunzi.Lib.Hku.Hk/Hkuto/Record/B3205189X*](Http://Sunzi.Lib.Hku.Hk/Hkuto/Record/B3205189X).

Ujvari, B., & Madsen, T. (2009). Short telomeres in hatchling snakes: Erythrocyte telomere dynamics and longevity in tropical pythons. *PLOS ONE*, *4*(10), e7493. <https://doi.org/10.1371/journal.pone.0007493>

Virginia Herpetological Society. (n.d.). Retrieved May 24, 2024, from http://www.virginiaherpetologicalsociety.com

Xu, M., Wu, X.-B., Yan, P., & Zhu, H. (2009). Telomere length shortens with age in Chinese alligators (*Alligator sinensis*). *Journal of Applied Animal Research*, *36*(1), 109–112. <https://doi.org/10.1080/09712119.2009.9707042>

Young, R. C., Kitaysky, A. S., Haussmann, M. F., Descamps, S., Orben, R. A., Elliott, K. H., & Gaston, A. J. (2013). Age, Sex, and Telomere Dynamics in a Long-Lived Seabird with Male-Biased Parental Care. *PLOS ONE*, *8*(9), e74931. <https://doi.org/10.1371/journal.pone.0074931>

Zou, Y., Yi, X., Wright, W. E., & Shay, J. W. (2002). Human telomerase can immortalize Indian muntjac cells. *Experimental Cell Research*, *281*(1), 63–76. <https://doi.org/10.1006/excr.2002.5645>
