## Supporting information for "Why are telomeres the length that they are? Insight from a phylogenetic comparative analysis"

**Addressing potential confounding variables**

*Methodology*

Given that telomere restriction fragment analysis (TRF) can be performed with or without denaturing DNA, we designated studies as denaturing, non-denaturing, a combination of denaturing and non-denaturing for meta-analyses, or unclear. We marked studies where methodology was unclear or where they used a combination of methodologies as “N/A” when testing for an effect of methodology. We then constructed a gls model with telomere length as the predictor variable and telomre length as the response variable. As methodology did not have a significant effect on any of our models (t_113_ = 0.51, p = 0.61), data from all methodologies listed above was included in our analyses (see supplied Dryad files for detailed results).

*Tissue type*

To assess the effect of tissue on telomere length measurements, we constructed a gls model and included tissue type as the predictor variable and telomere length as the response variable. We marked tissue type as “N/A” for studies that utilized multiple tissues when testing for an effect of tissue type to avoid potentially confounding effects. We found no significant effect of tissue type on telomere length (t_110_ < 0.247, p > 0.81 for all tissues). Additionally, a post-hoc pairwise comparison using the *emmeans* package found no significant differences in telomere length based on tissue type. As such, we included data from all tissue types in our analyses involving telomere length, lifespan, body mass, and baseline metabolism. Additionally, because measuring telomere length and telomerase activity in different tissues could lead to erroneous conclusions, we included only studies which measured both variables from the same tissue type when analyzing the correlated evolution of telomere length and telomerase activity.

*Standardization of telomere length measurements between studies*

When possible, we analyzed data that was analyzed in previous meta-analyses, reviews and studies that have collected data on telomere length from multiple sources (Gomes et al. 2011, Criscuolo et al. 2021, Pepke and Eisenberg 2022). This covers the vast majority of mammal and bird species. We applied the same correction factor to data from Seluanov et al. (2007) that precious analyses used (Pepke and Eisenberg 2022). As data on telomere length in fishes and non-avian archelisaurs and a handful of birds and mammals has not previously been analyzed in a comparative study, we were unable to apply a correction factor. However, given that telomere length measurements not including qPCR are highly repeatable, and that our study provides valuable insight, we included this data in our analyses (Lai et al. 2018, Kärkkäinen et al. 2022).

*Domestication*

Given that previous literature has demonstrated that the relationship between lifespan and telomere length is significantly impacted by the inclusion or exclusion of domesticated species, we did not include domesticated mammals in our analyses. The species excluded were the chicken (*Gallus gallus)*, pig (*Sus scrofa*), cow (*Bos taurus*), dog (*Canis lupus familiaris*), horse (*Equus caballus*), house mouse (*Mus musculus*), Norway rat (*Rattus norvegicus*), cat (*Felis catus*), sheep (*Ovis aries*), dromedary camel (*Camelus dromedarius),* and goat (*Capra hircus*).

*The calculation of telomere length and average adult telomere length*

Telomere rate of change differs drastically among species, and telomere attrition and lengthening occurs at different times in different species (Whittemore et al., 2019). Given this, standardizing adult telomere length among species is a difficult task, and a general approximation based on several individuals across a range of ages within a species may be the best indicator of adult telomere length. We excluded studies with measurements from extremely young or old individuals. For the majority of birds and mammals, we utilized the average adult telomere length values reported in previous analyses (Gomes et al. 2011, Criscuolo et al. 2021, Pepke and Eisenberg 2022). For species not included in these analyses, we used the reported average telomere length when available. For studies that did not report this, we calculated the average telomere length by adding up the measured telomere length for each individual and dividing by the number of individuals. For studies that reported telomere length as a range between two values, we recorded telomere length as the median value within this range.

**Alterations to phylogeny**

Species that lacked easily-accessible phylogenetic information were replaced with closely related species and, when possible, congeners in order to increase our sample size. The following alterations were made:

*Urolophus paucimaculatus* replaced with *Urogymnus asperrimus
Ailurus fulgens* replaced with *Ailurus
Giraffa camelopardalis* replaced with *Giraffa reticula
Saimiri sciureus* replaced with *Saimiri boliviensis
Fregata minor* replaced with *Fregata
Larus crassirostris* replaced with *Larus marinus
Trachemys scripta* replaced with *Trachemys dorbigni
Elephantulus rufescens* replaced with *Elephantulus brachyrhynchus*
